# Loss of ACTA1 leads to delayed γ-AChR / ε-AChR switch in skeletal muscle in mice

**DOI:** 10.64898/2026.05.22.727267

**Authors:** Yun Liu, Qiaohong Ye, Weichun Lin

## Abstract

Skeletal muscle actin forms the core structural component of thin filaments, which interact with thick filaments to generate contractile force. In addition to force production, the character of muscle contraction activity itself is thought to provide mechanical cues that influence synaptic development and maturation. In mouse skeletal muscle there is an early post-natal switch from embryonic forms of actin to the adult isoform, ACTA1, which increases both filament stability and force production. Newborn mice deficient for ACTA1 (*Acta1^−/−^*), although initially able to breath, move and suckle, develop profound muscle weakness and die during the early neonatal period, despite a compensatory, increase in expression of embryonic actins. We took advantage of this to better understand the response of the neuromuscular junction (NMJ) to a disruption in contractility and activity-dependent signaling during development. Morphological analyses of the diaphragm in *Acta1^−/−^* mice revealed that the patterning and formation of the NMJ proceed normally through postnatal day 5 (P5), the day at which pups begin to die. Short-term synaptic plasticity, assessed as the endplate potential (EPP) response to paired-pulse stimulation, was also unchanged, indicating normal presynaptic release of neurotransmitters. In contrast, electrophysiological recordings demonstrated significantly prolonged rise and decay kinetics of miniature and evoked endplate potentials, indicating altered postsynaptic receptor properties. Consistent with these functional changes, quantitative real-time PCR showed a reduced ratio of ε- to γ-acetylcholine receptor (AChR) subunit mRNA, reflecting a delay in the developmental switch from embryonic γ-containing to adult ε-containing AChRs. Together, these findings indicate that α-skeletal actin is dispensable for early NMJ morphogenesis but is required for timely postsynaptic receptor maturation, demonstrating a critical role for muscle contractile activity in coordinating synaptic development at the NMJ.

**Highlights:**

1. Skeletal muscle α-actin (ACTA1) is the principal structural component of thin filaments and a key determinant of contractile activity.
2. Using *Acta1*^−/−^ mice, we show that NMJ patterning and early morphogenesis occur normally despite severe impairment in muscle contractility.
3. Electrophysiological analysis of the NMJ shows that presynaptic function remains intact, as evidenced by normal paired-pulse responses. In contrast, postsynaptic maturation is disrupted, with prolonged endplate potential kinetics indicating altered AChR function.
4. This defect is associated with a delayed γ- to ε-AChR subunit switch, a key step in postnatal NMJ maturation. These findings identify ACTA1-dependent contractile activity plays a critical role in timely postsynaptic receptor maturation.

## Introduction

Actin is a highly conserved protein present in all eukaryotic cells and is essential for a wide range of fundamental cellular processes, including cell motility, cell division, intracellular transport, and muscle contraction. The mammalian actin family comprises six closely related isoforms, including two non-muscle (cytoplasmic β-actin and γ-actin) and four muscle actins (α-skeletal, α-cardiac, α-smooth, and γ-smooth actin) (Kashina, 2020; Perrin and Ervasti, 2010; Vandekerckhove and Weber, 1978). Muscle actin isoforms are expressed in highly regulated, developmental- and tissue-specific patterns. For example, during embryonic development, both α-vascular smooth muscle actin and α-cardiac actin are the predominant isoforms transiently expressed in skeletal muscle during embryonic development; however, α-skeletal actin (ACTA1) becomes the predominant isoform around birth and remains the principal component of thin filaments in adult skeletal muscle (McHugh et al., 1991; Vandekerckhove et al., 1986).

ACTA1 encodes skeletal muscle α-actin, the core component of the thin filament that interacts with myosin to drive muscle contraction. In addition to force generation, contractile activity produces activity-dependent and mechanical signals that regulate neuromuscular development, contributing to motor neuron refinement and synaptic stabilization at the NMJ (Olson and Nordheim, 2010). In humans, mutations in ACTA1 cause a spectrum of congenital myopathies including nemaline myopathy and congenital fiber-type disproportion (Costa et al., 2004; Ilkovski et al., 2001; Laing et al., 2004; Nowak et al., 2007; Nowak et al., 1999). In mice, complete loss of ACTA1 (*Acta1^−/−^*) results in neonatal lethality, accompanied by severely reduced muscle force generation and malnutrition (Crawford et al., 2002). Transgenic expression of cardiac (fetal) actin in the skeletal muscle of *Acta1^−/−^* mice can rescue postnatal lethality but results in a 25% reduction in the force of muscle contraction (Ochala et al., 2013). Thus, the skeletal muscle at birth in *Acta1^−/−^* mice is functional but its contractile properties differ from those of adult skeletal muscle. Unable to switch to adult ACTA1 as embryonic forms of actin are ablated, the muscle progresses to a profound contractile dysfunction. We hypothesized that loss of muscle contractility also disrupts activity-dependent signals required for proper NMJ maturation. In particular, impaired force generation could alter postsynaptic receptor stabilization, AChR subunit composition, or retrograde signaling to motor nerve terminals. We therefore performed morphological and electrophysiological analyses of *Acta1^−/−^*mice to define the role of skeletal muscle actin in neuromuscular development and early postnatal synaptic function.

We found that NMJ formation and differentiation during embryonic and neonatal stages occur normally in the absence of skeletal actin. However, the kinetics of both spontaneous and evoked endplate potentials is prolonged, indicating altered postsynaptic AChR channel properties. Furthermore, quantitative real-time PCR revealed a reduced ratio of AChR ε-subunit to γ-subunit mRNA in mutant muscle at postnatal day 5 (P5). Together, these findings indicate that α-skeletal actin is dispensable for NMJ formation and early morphological maturation but is required for the timely developmental transition of AChRs from the embryonic to the adult form.

## Results

### NMJ formation and the onset of synapse elimination are normal in *Acta1^−/−^* mice

Mice lacking skeletal muscle actin (*Acta1^−/−^*) die during the neonatal period and exhibit severe muscle weakness, growth retardation, and malnutrition (Crawford et al., 2002). Because effective neuromuscular transmission is essential for postnatal motor behaviors such as feeding, we first asked whether loss of skeletal muscle actin disrupts NMJ development. *Acta1^+/−^* mice were intercrossed to generate *Acta1^−/−^* offspring. Within 2-3 days after birth, *Acta1^−/−^* mice were phenotypically indistinguishable from their wild-type or heterozygous littermates. However, starting from postnatal day 3 (P3), mutant mice exhibited smaller body size and less motility. Most *Acta1^−/−^* mice died between P5 to P9 and no mutant mice survived beyond P10. These observations are consistent with previous reports (Crawford et al., 2002). Due to the early neonatal lethality in *Acta1^−/−^*mice, we focused our analyses on two stages – embryonic day 18.5 (E18.5) and postnatal day 5 (P5), a stage prior to the onset of postnatal lethality.

Embryonic diaphragm muscles (E18.5) were double-labeled with Texas-Red conjugated *α*-bungarotoxin for AChRs and antibodies against syntaxin 1 for nerves. *Acta1^−/−^* and *Acta1^+/+^* diaphragm muscles exhibited a similar innervation pattern (Fig. 1A). High power images show presynaptic nerve terminals were juxtaposed to postsynaptic AChRs (Fig. 1B). These results indicate that initial NMJ formation occur normally in the absence of skeletal muscle actin.

**Figure 1.**
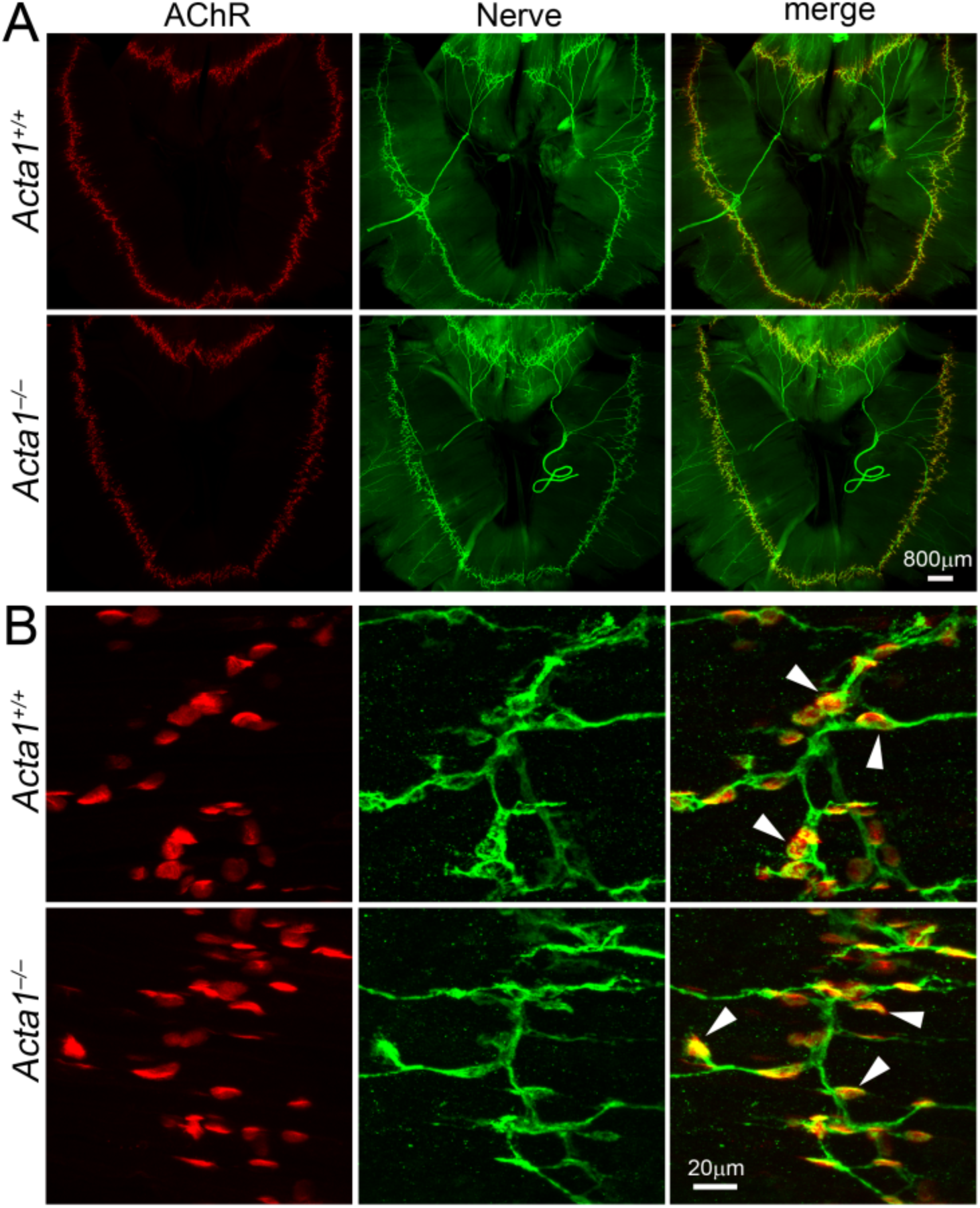
Innervation pattern and NMJ formation are normal in *Acta1^−/−^* mice at embryonic day E18.5. **A.** Whole-mount diaphragm muscles from E18.5 *Acta1^+/+^*and *Acta1^−/−^* embryos were double-labeled with Texas-Red conjugated *α*-bungarotoxin for AChRs (red) and antibodies against syntaxin 1 for nerves (green). *Acta1^−/−^* and *Acta1^+/+^* embryos exhibited similar innervation pattern in the diaphragm muscles. **B.** High power view of NMJs in **A**. Presynaptic nerve terminals (green) were aligned with postsynaptic AChRs (red) in both mutant and wildtype embryos (arrowheads).

At P5, the nerve terminals were also juxtaposed to postsynaptic AChR clusters in both control and mutant muscles (Fig. 2A). Furthermore, control and mutant neuromuscular junctions showed similar neonatal morphology of AChR clusters, characterized by oval plaques with homogenous *α*-bungarotoxin staining. To further assess postsynaptic differentiation, we examined the localization of acetylcholinesterase (AChE), a hallmark of mature postsynaptic membranes (Rotundo, 2003). We double-labeled diaphragm muscles with antibodies against AChE and Texas-Red conjugated *α*-bungarotoxin. We found that AChE staining was co-localized with *α*-bungarotoxin labeled AChR clusters in both control and mutant muscles (Fig. 2 B). Additionally, quantitative measurement of the AChR clusters (endplates) revealed similar sizes in control and mutant mice (Fig. 2 C). These results suggest that postnatal development and differentiation of the NMJ proceed normally in *Acta1^−/−^* mice.

**Figure 2.**
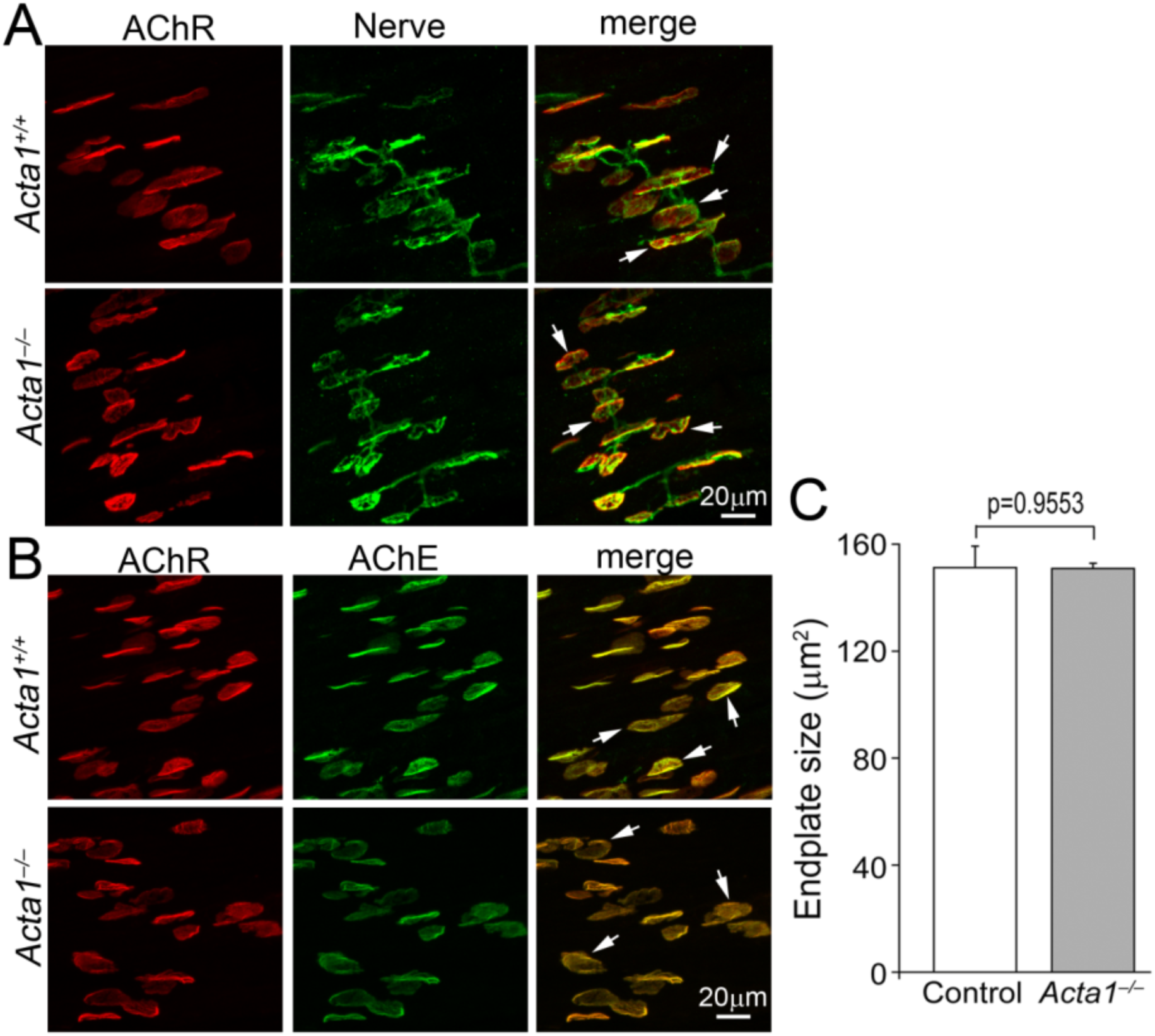
Postnatal NMJ morphology is normal in *Acta1^−/−^* mice at P5. **A.** Whole-mount diaphragm muscles from P5 *Acta1^+/−^* and *Acta1^−/−^*mice were double-labeled with Texas-Red conjugated *α*-bungarotoxin for AChRs (red) and antibodies against syntaxin 1 for nerves (green). In both control and mutant mice, anti-syntaxin 1 antibodies labeled both presynaptic nerves and nerve terminals. Nerve terminals were juxtaposed to postsynaptic AChRs (arrows). **B.** Whole-mount diaphragm muscles from P5 *Acta1^+/+^* and *Acta1^−/−^* mice were double-labeled with antibodies against acetylcholinesterase (AChE) (green) and Texas-Red conjugated *α*-bungarotoxin for AChRs (red). In both control and mutant muscles, *α*-bgt labeled AChR clusters were also co-labeled with anti-AChE antibodies (arrows). **C.** The size of endplates in P5 diaphragm muscles. No significant difference in endplate size was found between mutant (150.8 ± 0.2 μm, N=3 mice, n=161 endplates) and control mice (151.2 ± 8 μm, N=3, n=144), P=0.9553, Student’s *t*-test.

During early postnatal development, individual muscle fibers are initially innervated by multiple motor axons. This process is followed by activity-dependent synapse elimination that refines innervation to a single axon per endplate by 2 weeks postnatally (Sanes and Lichtman, 1999; Tapia et al., 2012; Thompson, 1985). To determine whether the deficiency of skeletal actin affected postnatal synapse elimination at the NMJ, triangularis sterni (TS) muscles at P5 were double labeled with antibodies against neurofilament-150 and synaptotagmin 2 to visualize axons and nerve terminals and with Texas-Red conjugated *α*-bungarotoxin to visualize AChRs. High-power confocal images were collected for quantitative analyses (Fig. 3A). In mutant TS muscles, the percentages of poly-innervated and mono-innervated NMJs were 75.59 ± 3.96 and 24.41 ± 3.96 respectively. These results were comparable to the results obtained from control TS muscles (poly-innervation 70.65 ± 5.66 and mono-innervation 29.34 ± 5.66) (Fig. 3B). Although the early lethality of *Acta1^−/−^*mice precludes examination of later stages of synapse elimination, these results indicate that the initial postnatal phase of synaptic pruning proceeds normally in the absence of skeletal muscle actin.

**Figure 3.**
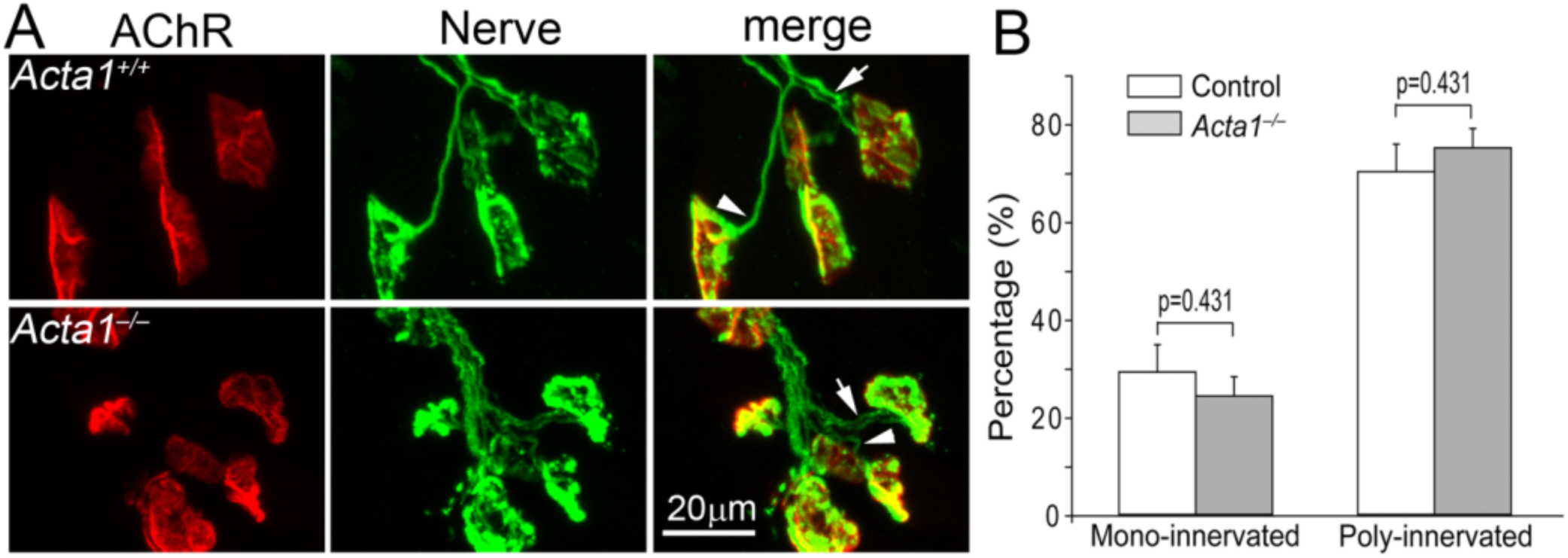
Early synapse elimination is not altered in *Acta1^−/−^* mice at P5. **A.** Whole-mount triangularis sterni (TS) muscles from P5 *Acta1^+/−^* and *Acta1^−/−^* mice were double-labeled with antibodies against neurofilament and synaptotagmin 2 for axons and nerve terminals (green) and Texas-Red conjugated *α*-bungarotoxin for AChRs (red). Arrows indicate the axons that poly-innervated the NMJ. Arrowheads indicate the axons that mono-innervated the NMJ. **B.** Quantification of mono-and poly-innervation in TS muscle. In mutant muscles, the percentages for mono- and poly-innervations were 24.41±3.96 and 75.59±3.96 respectively, which were comparable to those of the control muscles (29.34±5.66 and 70.65±5.66 respectively), P=0.431, Student’s *t*-test.

Together, these findings demonstrate that skeletal muscle actin is not required for embryonic NMJ formation, early postnatal differentiation, or the onset of synapse elimination.

### The time course of miniature and evoked endplate potentials are prolonged in *Acta1^−/−^* mice

Although NMJ morphology appeared normal in *Acta1^−/−^* mice, profound muscle weakness raised the possibility that synaptic function might be altered. We performed intracellular recordings on phrenic nerve-diaphragm muscle preparations from *Acta1^−/−^*and their littermate control (*Acta1^+/+^* or *Acta1^+/−^*) mice at P5. We first examined spontaneous transmitter release at the NMJ by recording miniature endplate potentials (mEPPs) which represent muscle membrane depolarization induced by the release of single synaptic vesicles (Fig. 4A). The frequency and amplitude of the mEPP in mutant muscles were comparable to those of the control (Fig. 4C, 4D). However, we noticed that compared to control, the mEPPs in mutants exhibited elongated event duration (overlaid traces in Fig. 4B). The analyses of mEPP’s time course showed marked increase in rise time and decay time in mutant muscles (Fig. 4E, 4F, 4G).

**Figure 4.**
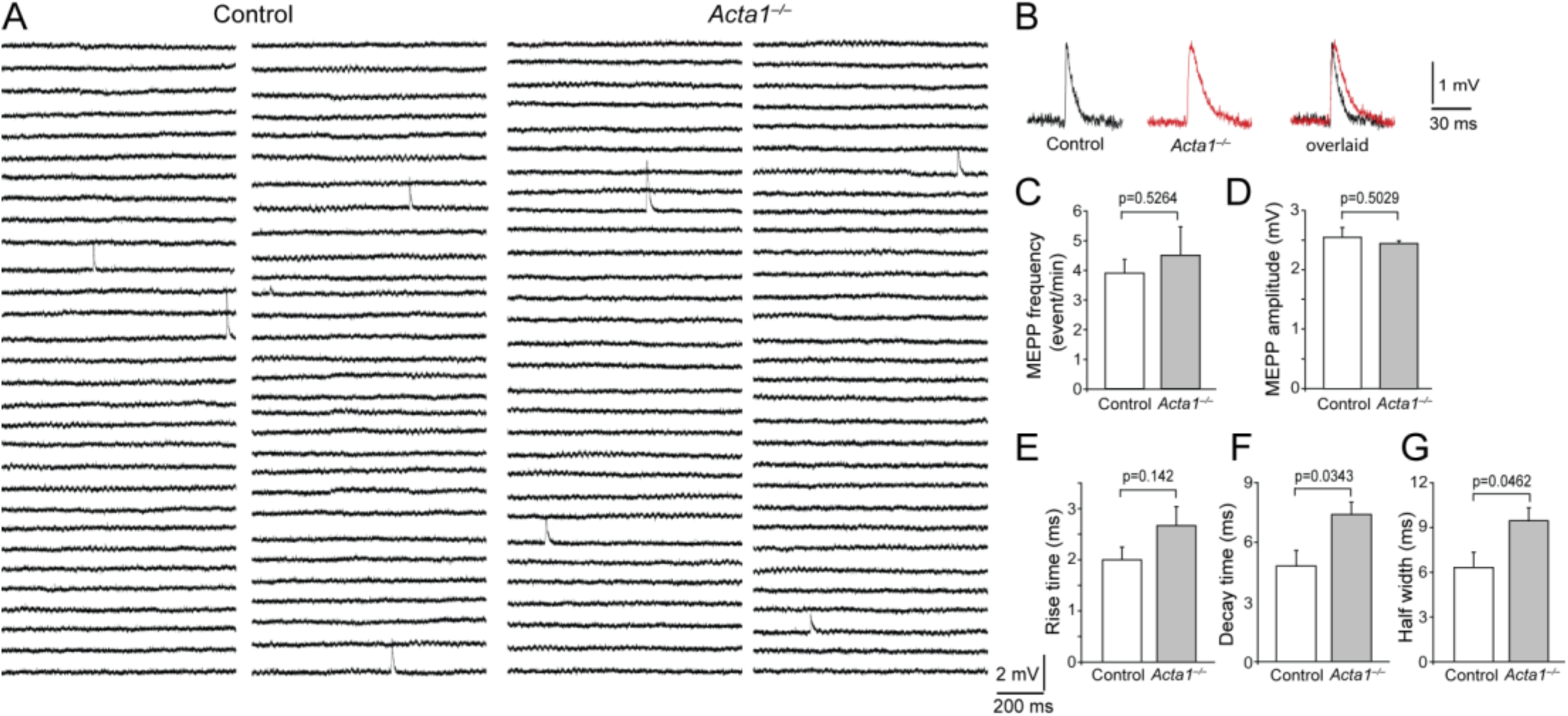
Miniature endplate potentials are prolonged in *Acta1^−/−^*muscles at P5. **A.** Sample traces of 1-minute continuous recording of mEPPs from control and *Acta1^−/−^*diaphragm muscles at P5. **B.** Traces of individual mEPPs recorded from control (black trace) and *Acta1^−/−^* (red trace) muscle. mEPP in mutant muscle (red trace) shows longer duration when overlaid (right panel). **C, D.** Quantification of mEPP frequency and amplitude. No significant changes were detected in mutant mice. mEPP frequency: control, 3.93 ± 0.46 events/min, N = 3 mice, n = 35 cells; mutant: 4.53 ± 0.97 events/min, N = 3, n = 40. mEPP amplitude: control, 2.57 ± 0.17 mV; mutant, 2.47 ± 0.05 mV. **E-G.** Quantification of rise time (10% - 90%) (E), decay time (100% - 50%) (F) and half width (G) of mEPPs. Compared to those of the control mice, the three parameters of time course significantly increased in mutant mice: rise time in mutant mice increased by 34% from the control, 2 ± 0.25 ms to 2.67 ± 0.37 ms; decay time increased by 54% from 4.81 ± 0.79 ms to 7.39 ± 0.62 ms; half width increased by 50% from 6.32 ± 1.02 ms to 9.46 ± 0.88 ms.

To determine evoked neurotransmitter release at the NMJs, we applied supra threshold stimuli to the phrenic nerve and recorded endplate potentials (EPPs) in the muscle (Fig. 5A). The amplitude and quantal content of EPPs were similar between mutant and control mice (Fig. 5B, 5C); in contrast, the duration of the EPPs was significantly longer in mutant mice (overlaid traces in Fig. 5A). The analyses of time course revealed significant increases in rise time and decay time in mutant mice compared to control (Fig. 5D, 5E, 5F). These changes in time course observed in both spontaneous and evoked responses suggest that loss of skeletal muscle actin affects postsynaptic response kinetics rather than presynaptic transmitter release.

**Figure 5.**
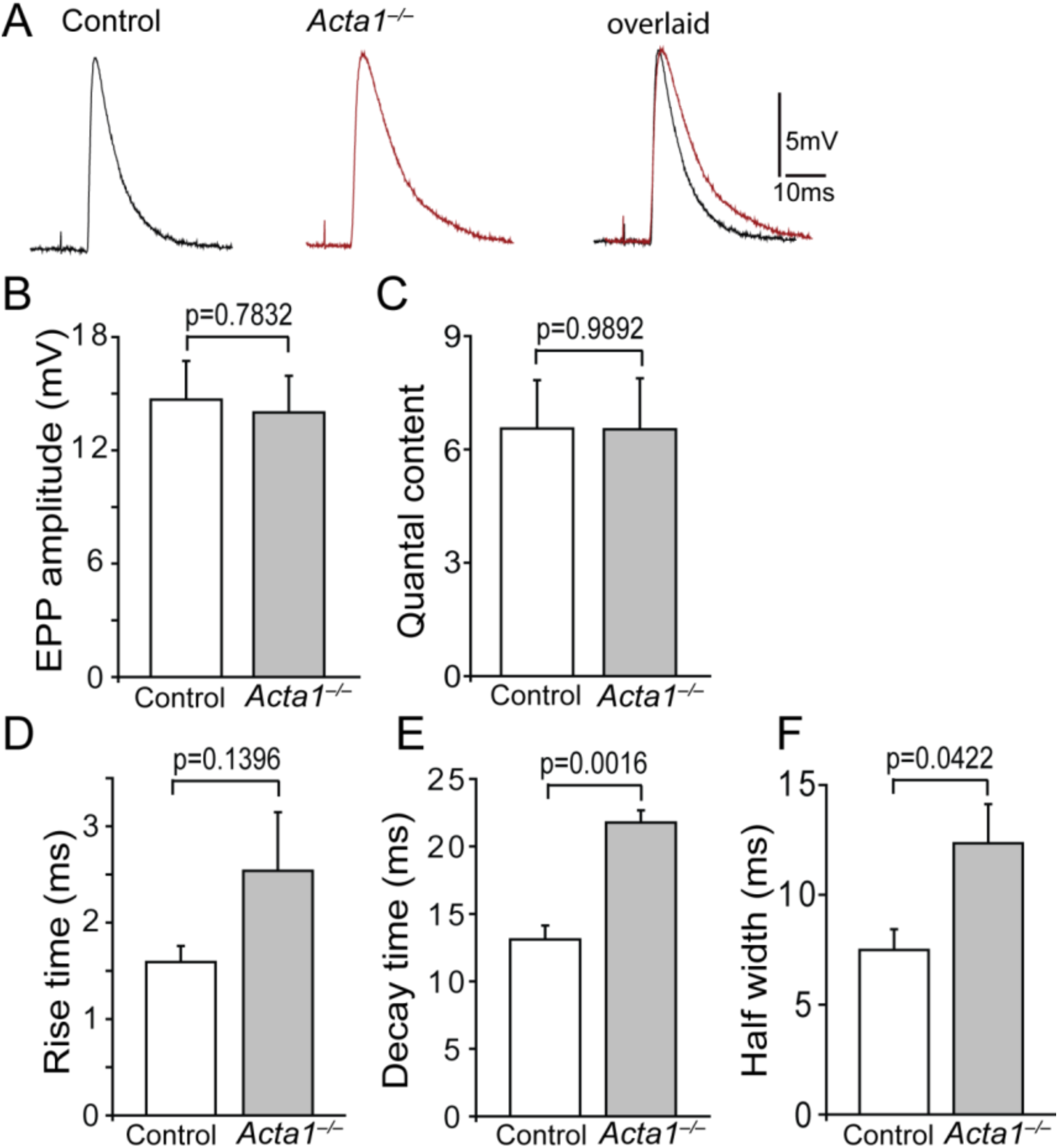
The time course of evoked endplate potential is prolonged in *Acta1^−/−^*muscles at P5. **A.** Sample traces of evoked endplate potentials (EPPs) recorded from control and *Acta1^−/−^* mice at P5. The right panel shows overlaid control and mutant EPP traces. **B.** Quantification of EPP amplitude. No significant difference was observed between control (14.68 ± 2.05 mV, N = 3 mice, n = 29 cells) and mutant mice (14 ± 1.94 mV, N = 3, n = 34). **C.** Quantification of quantal content. No significant difference was observed between control (6.56 ± 1.28) and mutant mice (6.53 ± 1.35). **D-F.** Quantification of rise time (10% - 90%) (D), decay time (90% - 10%) (E) and half width (F) shows significant increases in mutant mice, compared to that of the control: rise time in mutant mice increased by 60% from the control, 1.59 ± 0.17 ms to 2.54 ± 0.61 ms; decay time increased by 66% from 13.11 ± 1.03 ms to 21.76 ± 0.92 ms; half width increased by 65% from 7.49 ± 0.95 ms to 12.34 ± 1.78 ms.

### Short-term synaptic plasticity is preserved at the NMJs in *Acta1^−/−^* mice

Short-term synaptic plasticity refers to the temporary increase (facilitation) or decrease (depression) in synaptic strength in response to repeated stimulation within a timescale of milliseconds to a few minutes. Paired-pulse stimulation with various stimulation intervals (150ms, 100ms and 50 ms) was applied to the phrenic nerve and the EPPs were recorded in the muscle (Fig. 6A). The amplitude ratio of the second to first EPP was calculated. The ratio of EPP(2) / EPP(1) indicates synaptic facilitation when the value is more than 1. In control mice, the ratios were 1.2 ± 0.04 (150ms), 1.25 ± 0.05 (100ms) and 1.43 ± 0.05 (50ms); in mutant mice, the ratios were 1.12 ± 0.03 (150ms), 1.19 ± 0.04 (100ms) and 1.4 ± 0.08 (50ms). There were no significant differences between mutant and control mice (Fig. 6B). Furthermore, we applied 10 Hz repetitive stimulation to the nerve and recorded EPPs in the muscle (Fig. 6C). In control mice, the NMJ exhibited synaptic facilitation throughout the stimulation, which was revealed by the ratios of the EPP(n) to EPP(1) that were more than 1. A similar pattern was observed in mutant mice (Fig. 6D). These results indicate that short-term synaptic plasticity at the NMJ is similar between *Acta1^−/−^* and control mice.

**Figure 6.**
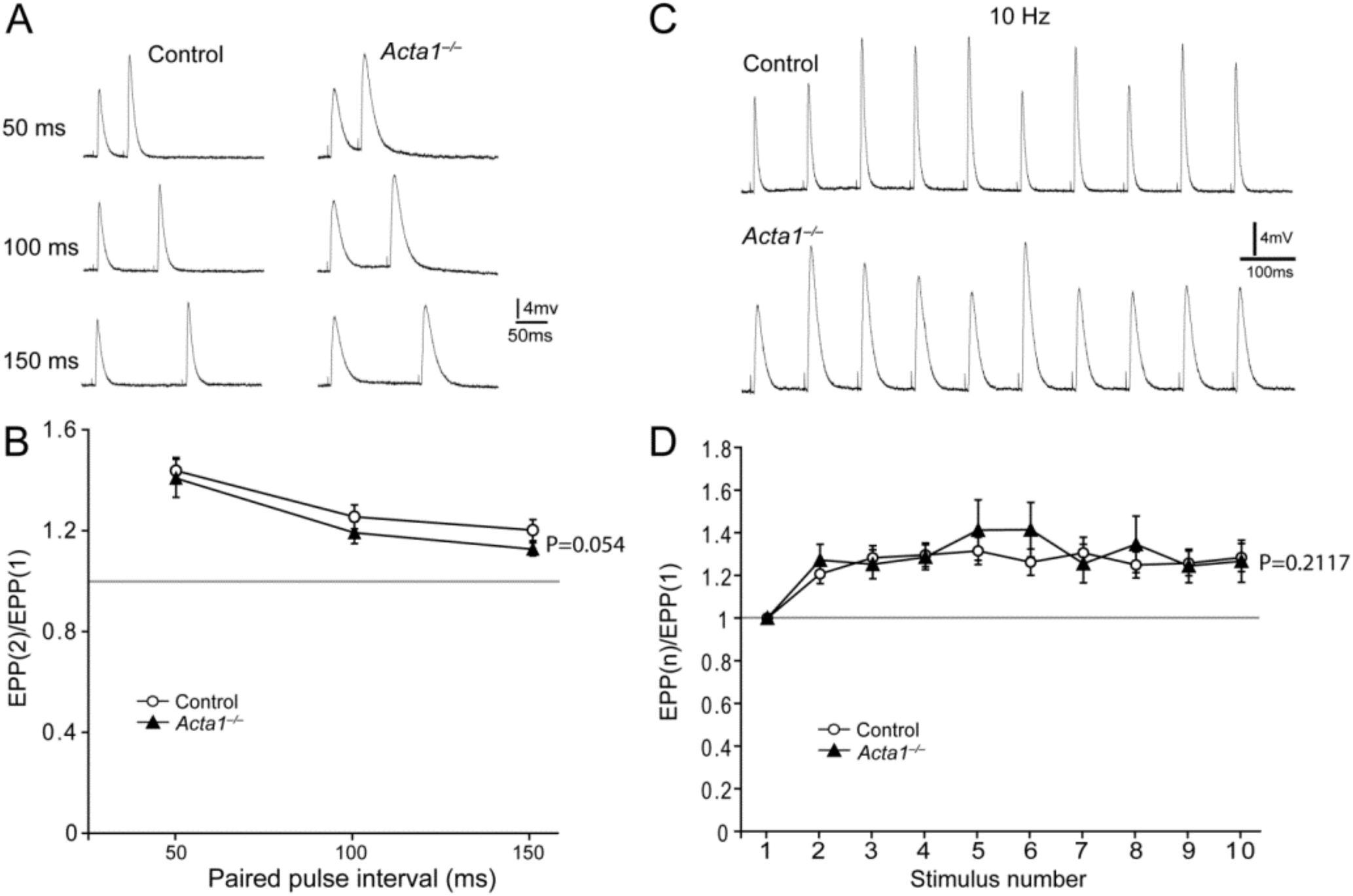
Short-term synaptic plasticity at the NMJ is normal in *Acta1^−/−^* mice at P5. **A.** Sample traces of EPPs responding to pair-pulse stimulation at various time intervals (50-150ms). **B.** Paired pulse ratios of EPP(2) to EPP(1). No significant difference in paired-pulse facilitation was detected between control and mutant mice. **C.** Sample traces of EPPs recorded during 10 Hz repetitive stimulation. **D.** The ratios of EPP(n) to EPP(1) show that both control and mutant neuromuscular junctions exhibited similar EPP facilitation responding to 10 Hz nerve stimulation and there was no difference between control and mutant mice. Control: N = 3 mice, n = 29 cells; *Acta1^−/−^*: N = 3 mice, n = 34 cells.

### Delayed γ-AChR / ε-AChR switch in *Acta1^−/−^* mice

Electrophysiological analyses revealed normal mEPP frequency and amplitude, EPP amplitude, and short-term synaptic plasticity, indicating that presynaptic neurotransmitter release is largely unaffected by the loss of skeletal muscle actin. In contrast, the prolonged kinetics and half-width of mEPPs and EPPs in *Acta1^−/−^* mice suggest significant alterations in postsynaptic AChR channel properties. AChRs are ligand (acetylcholine)-gated ion channels located in the postsynaptic muscle membrane. During postnatal development, AChR maturation involves the replacement of the γ subunit with the ε subunit, a transition that produces distinct changes in AChR channel properties.

At P5, both γ- and ε-containing AChRs are co-expressed in the same individual endplates in mouse diaphragm muscles (Missias et al., 1996; Yumoto et al., 2005). We therefore hypothesized that the prolonged synaptic responses observed in *Acta1^−/−^* mice might reflect a delay in this developmental AChR subunit switch. To test this, we quantified the mRNA expression levels of AChR subunits in diaphragm muscles at P5 using RT-qPCR (Fig. 7A). We examined α, β, δ, γ, and ε subunits and used housekeeping gene *GAPDH* as reference control. The results show that the mRNA levels of AChR subunits α, β, γ, and ε were significantly reduced in mutant mice, whereas expression of the δ subunit was not affected (Fig. 7A).

**Figure 7.**
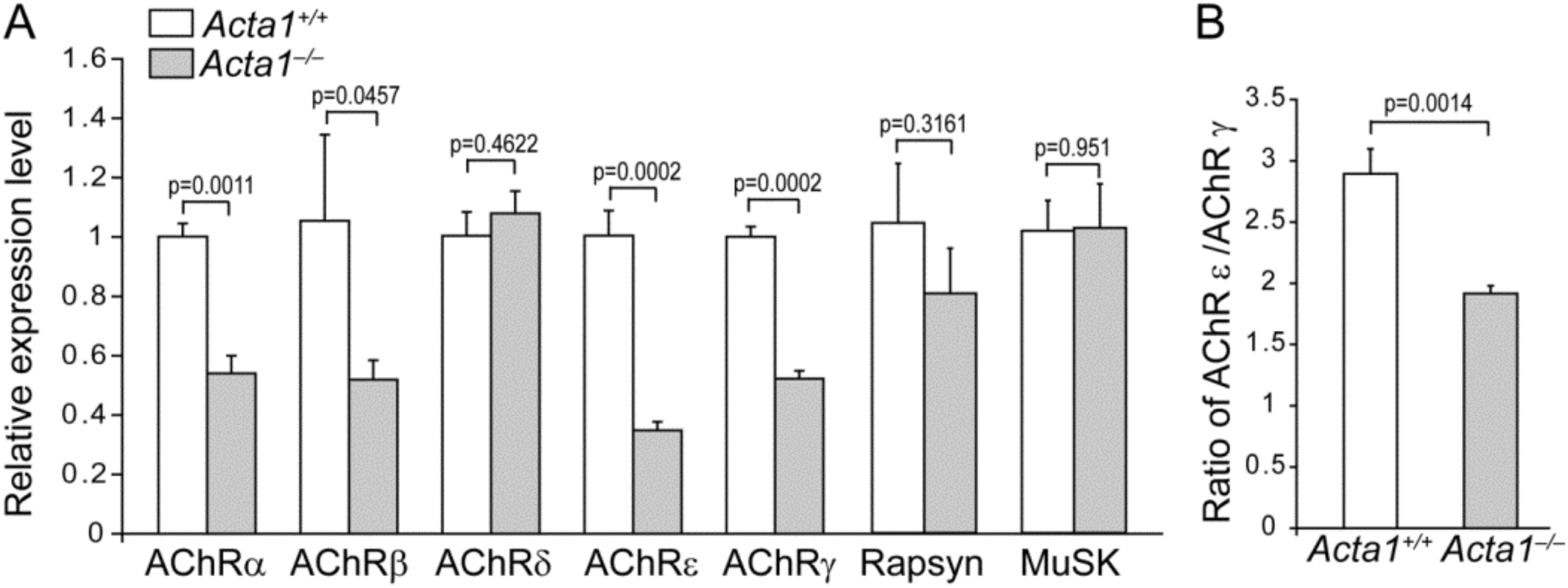
Gene expression of adult isoforms of AChR subunits is delayed in *Acta1^−/−^* mice. **A.** Relative mRNA levels of AChR subunits α, β, δ, ε, γ in diaphragm muscles of *Acta1^+/+^* and *Acta1^−/−^*mice at P5. Compared to those of *Acta1^+/+^* mice, the relative mRNA levels of AChR subunits α (0.54±0.06), β (0.52±0.07), ε (0.35±0.03), and γ (0.52±0.03) were significantly reduced in *Acta1^−/−^*mice. The mRNA levels of AChR subunit δ, Rapsyn and MuSK were similar between *Acta1^+/+^* and *Acta1^−/−^* mice. **B.** The ratio of AChR subunit ε / γ was significantly reduced in *Acta1^−/−^* mice (1.92±0.06) compared to that of *Acta^+/+^* mice (2.89±0.2). N = 4 mice per genotype.

Notably, the ratio of ε- to γ-subunit was 2.89 in wildtype, suggesting AChRε was more abundant than AChRγ at P5 in wildtype muscles. In contrast, the ratio of ε / γ was 1.92 in mutant mice, significantly lower than the ratio of 2.89 in wildtype muscles (P = 0.0014), indicating a marked reduction in ε subunit expression compared to that of wildtype mice (Fig. 7B). These results demonstrate that the postnatal transition from embryonic γ-containing AChRs to adult ε-containing receptors is delayed in the absence of skeletal muscle actin.

We also determined the mRNA levels of two critical proteins in the postsynaptic muscle membrane, rapsyn and MuSK that are required for the development and maintenance of NMJs (Burden et al., 2003; DeChiara et al., 1996; Gautam et al., 1999; Valenzuela et al., 1995; Xie et al., 2023). We found that the mRNA levels of Rapsyn and MuSK were similar between WT and *Acta1^−/−^* muscles (Fig.7 A), consistent with the morphological observations on mutant diaphragms at E18.5 and P5 which show normal development and formation of NMJs.

## Discussion

Proper NMJ development, maturation, and maintenance depend on coordinated muscle activity and intact muscle structure. Disruption of muscle function, including that caused by muscle disorders and pathological conditions, can impair NMJ development, postnatal maturation, and synaptic stability (Chen et al., 2011; Gonzalez-Freire et al., 2014; Liu and Lin, 2023; Lovering et al., 2020; Rudolf et al., 2014; Theroux et al., 2008; van der Pijl et al., 2016). For example, a dominant ACTA1 (H40Y) mutation causes severe muscle weakness and nemaline myopathy in both human patients and a corresponding *Acta1*(H40Y) knock-in mouse model (Nguyen et al., 2011; Nowak et al., 1999). Consistent with these findings, we previously demonstrated pronounced structural and functional defects at the NMJ in this mouse model, indicating that perturbation of skeletal muscle actin disrupts NMJ structure, function and maintenance (Liu and Lin, 2023). However, in this case it was not possible to tell whether changes in the NMJ were due to physical disruption by nemaline bodies or a response to contractile deficiency due to reduced actin function. Thus, we turned to the analysis of animals lacking α-skeletal actin (ACTA1).

Skeletal muscle contraction is initiated by excitation–contraction (E–C) coupling, which begins when acetylcholine (ACh) released from presynaptic motor nerve terminals binds to postsynaptic ACh receptors (AChRs) at the neuromuscular junction (NMJ). This triggers depolarization of the postsynaptic membrane, generating end-plate potentials and propagating action potentials along the muscle fiber. These action potentials activate L-type calcium channels, also known as dihydropyridine receptors (DHPRs), in the transverse (T-) tubules, which in turn activate ryanodine receptors (RyRs), leading to calcium release from the sarcoplasmic reticulum and subsequent muscle contraction.

In *Acta1^−/−^* mice, muscle force generation is severely impaired (Crawford et al., 2002), providing a model to assess the role of contraction in NMJ development. We find that NMJ patterning and initial formation proceed normally in *Acta1^−/−^* mice at both E18.5 and P5. These data indicate that muscle contraction per se is not required for the establishment of NMJ patterning or early synapse formation. These findings are consistent with our previous work showing that NMJ patterning depends on skeletal muscle DHPR function, but not on downstream E-C coupling (Chen et al., 2011). Importantly, the present study extends this conclusion by directly excluding a role for muscle contraction in regulating NMJ patterning and formation during development.

However, muscle contraction plays a critical role in NMJ maturation. We find that postnatal maturation of AChRs is delayed in *Acta1^−/−^*mice, as evidenced by a marked reduction in the ε- to γ-AChR subunit mRNA ratio. This results in the persistence of embryonic AChR channel properties into the postnatal period. These findings indicate that while skeletal muscle actin - and thus effective force generation - is dispensable for early NMJ assembly, it is required for the postnatal maturation of AChRs.

AChRs are pentameric ligand-gated ion channels that mediate synaptic transmission at the NMJ. During development, AChRs undergo a well-characterized subunit switch from the embryonic γ-containing form (α₂βγδ) to the adult ε-containing form (α₂βεδ) (γ-AChR / ε-AChR switch) (Duclert and Changeux, 1995; Li et al., 2024; Mishina et al., 1986). In mice, this transition occurs during the first two postnatal weeks and is a hallmark of NMJ maturation (Missias et al., 1996; Yumoto et al., 2005). Substitution of the γ subunit with ε confers distinct functional properties on the receptor. Incorporation of the ε subunit confers distinct biophysical properties, including reduced acetylcholine sensitivity, increased single-channel conductance, and shorter channel open times, enabling rapid and reliable neuromuscular transmission in mature muscle (Cetin et al., 2020; Fischbach and Schuetze, 1980; Kopta and Steinbach, 1994; Li et al., 2024; Mishina et al., 1986; Schuetze and Vicini, 1986).

The prolonged decay time and increased half width of mEPPs and EPPs observed in *Acta1^−/−^* mice suggest that the γ-to-ε AChR transition is delayed. This interpretation is supported by previous studies demonstrating that deletion of the AChR ε subunit results in prolonged decay of miniature endplate currents due to the persistence of γ-AChRs at adult endplates (Missias et al., 1997; Witzemann et al., 1996). Conversely, in mice lacking the AChR γ subunit, the half width of mEPPs is significantly reduced in E18.5 embryos because ε-containing AChRs become predominant (Liu et al., 2008; Liu et al., 2010). Together, these findings are consistent with our electrophysiological observations and indicate that embryonic AChRs persist longer in *Acta1^−/−^* muscle.

The γ-to-ε AChR switch is regulated transcriptionally within subsynaptic muscle nuclei. The gene transcription during synapse development is influenced by motor innervation, calcium influx, and muscle activity (Brenner et al., 1994; Cohen et al., 1997; Duclert and Changeux, 1995; Kues et al., 1995; Missias et al., 1996; Olson and Nordheim, 2010; Rimer et al., 1997). Our electrophysiological analyses indicate that presynaptic transmission is largely preserved at the NMJs in *Acta1^−/−^* mice, suggesting that presynaptic input is unlikely to account for the delayed AChR transition. Instead, reduced muscle contractile activity may play a key role. Although cardiac and smooth muscle actin isoforms are upregulated in *Acta1^−/−^* mice as a compensatory response (Crawford et al., 2002), this compensation does not fully restore contractile function. Consistent with this, myofibres from *Acta1^−/−^* mice rescued by transgenic expression of cardiac actin in skeletal muscle generate significantly less force than wild type myofibres (Ochala et al., 2013).

We postulate that reduced muscle contractility weakens the activity-dependent signaling pathways critical for the timely induction of AChR ε-subunit expression during postnatal synapse development. In addition, actin dynamics may directly influence transcriptional regulation of muscle genes through the actin-MRTF-SRF signal pathway (Olson and Nordheim, 2010). In this pathway, changes in actin polymerization regulate the nuclear localization of myocardin-related transcription factors (MRTFs), which in turn activate serum response factor (SRF)-dependent gene expression. In this view, skeletal muscle actin may contribute to postsynaptic gene regulation not only by supporting contractile activity but also by modulating cytoskeleton-dependent transcriptional signaling during synaptic maturation.

An alternative explanation of our findings is that the delayed AChR switch arises secondarily from growth retardation, malnutrition, or general developmental delay associated with early neonatal lethality in *Acta1^−/−^* mice. Distinguishing between primary and secondary mechanisms will require experimental strategies that prolong postnatal survival while selectively manipulating muscle contractile function. Such approaches will help determine whether muscle contractility directly controls the developmental timing of AChR subunit switching.

## Materials and methods

### Mice

Heterozygous *Acta1^+/−^* mice were obtained from Dr. Kristen Nowak (The University of Western Australia), who has maintained *Acta1^+/−^* mouse line that was previously generated by Dr. James Louis Lessard (Cincinnati Children’s Hospital Medical Center) (Crawford et al., 2002). Heterozygous *Acta1^+/−^*mice were viable and fertile, and were intercrossed to generate *Acta1^−/−^* offspring. Litters were monitored daily after birth. Consistent with previous reports (Crawford et al., 2002), *Acta1^−/−^* mice were born at the expected Mendelian ratio and appeared indistinguishable from their littermate controls (*Acta1^+/+^ and Acta1^+/−^*) until postnatal day 3 (P3). After P3, mutant mice exhibited reduced body size, diminished mobility, and increased mortality, with all *Acta1^−/−^* mice dying by postnatal day 10. Due to this neonatal lethality, all postnatal analyses were performed at P5. For embryonic studies, embryos were collected at embryonic day 18.5 (E18.5). To obtain mouse embryos, *Acta1^+/−^* mice were time-mated, and the day of vaginal plug detection was designated as E0.5. Pregnant females were anesthetized, and embryos were harvested by cesarean section at E18.5. All experimental protocols followed National Institutes of Health (NIH) guidelines and were approved by the University of Texas Southwestern Institutional Animal Care and Use Committee.

### Immunofluorescence staining

Whole mount immunofluorescence staining was carried out as previously described (Liu et al., 2008; Liu et al., 2025). Briefly, either diaphragm or triangularis sterni muscles from *Acta1^−/−^* and their littermate controls (*Acta1^+/+^* or *Acta1^+/−^*) at E18.5 or P5 were fixed in 2% paraformaldehyde in 0.1 M phosphate buffer (pH 7.3) overnight at 4^°^C. Samples were extensively washed with PBS and then incubated with Texas-Red conjugated α-bungarotoxin (α-bgt) (2 nM, Invitrogen, Carlsbad, California, USA) for 30 minutes at room temperature. Samples were then incubated with primary antibodies overnight at 4^°^C. The following polyclonal antibodies were used: neurofilament (NF150) (Chemicon, Temecula, CA, USA), synaptotagmin 2 (I735), syntaxin 1(I375) (generous gifts from Dr. Thomas Südhof, Stanford University School of Medicine, Palo Alto, CA, USA) (Pang et al., 2006), and acetylcholinesterase (AChE) (generous gifts from Dr. Palmer Taylor, UC San Diego, CA, USA). All primary antibodies were diluted by 1:1000 in antibody dilution buffer (500 mM NaCl, 0.01 M phosphate buffer, 3% BSA, 0.01% thimerosal and 1% Triton X-100). After extensive washes with PBST (1% Triton X-100 in PBS), muscle samples were incubated with fluorescein isothiocyanate (FITC)-conjugated goat anti-rabbit IgG (1:600, Jackson ImmunoResearch Laboratories, Inc., West Grove, PA, USA) overnight at 4^°^C. Muscle samples were then washed in PBST, followed by PBS, and mounted in Vectashield mounting medium (H-1000, Vector Laboratories, Inc., Burlingame, CA, USA). Images were acquired using a Zeiss LSM 880 confocal microscope. Confocal images captured at high magnification (63× oil /N.A. 1.4) were used to measure endplate area by using NIH ImageJ software. The endplate area was measured as the thresholded area labeled by Texas-Red conjugated α-bgt (Liu and Lin, 2023).

### Electrophysiology

Electrophysiological analyses was performed on 3 pairs of *Acta1^−/−^* and their littermate controls (*Acta^+/+^* or *Acta1^+/−^*) at P5. Intracellular recordings were carried out as previously described (Liu et al., 2011). Briefly, phrenic nerve-diaphragm muscles were dissected and mounted on a Sylgard coated dish, bathed in oxygenated (95% O_2_, 5% CO_2_) Ringer’s solution (136.8 mM NaCl, 5 mM KCl, 12 mM NaHCO_3_, 1 mM NaH_2_PO_4_, 1 mM MgCl_2_, 2 mM CaCl_2_, and 11 mM d-glucose, pH 7.3). Endplate regions were identified under a water-immersion objective (Olympus BX51WI) and impaled with glass microelectrodes (resistance 20–40 MΩ) filled with 2 M potassium citrate and 10 mM potassium chloride. To eliminate muscle action potentials and suppress muscle contractions, μ-conotoxin GIIIB (1 μM, Peptides International, Louisville, KY, USA) was added to the bath solution 30 minutes prior to recording. Supra threshold stimuli (2-5V, 0.1ms) were applied to the phrenic nerve via a glass suction electrode connected to an extracellular stimulator (SD9, Grass-Telefactor, West Warwick, RI). Signals were acquired with an intracellular amplifier (AxoClamp-2B) and digitized with Digidata 1332A (Molecular Devices, Sunnyvale, CA, USA). Data were analyzed with pClamp 10.7 (Molecular Devices) and Mini Analysis Program (Synaptosoft, Inc., Decatur, GA).

Amplitude measurements of mEPPs and EPPs were corrected for nonlinear summation using established methods (Liu and Lin, 2023; Rozas et al., 2011; Wood and Slater, 2001). The amplitudes of mEPPs and EPPs were normalized to −75 mV by using the formula EPP_normalized_ = EPP × (-75/V_m_) where V_m_ was the measured resting membrane potential (Rozas et al., 2011). The EPP_normalized_ was then corrected for non-linear summation by using the formula EPP’ = EPP_normalized_ / [1 – *f* (EPP_normalized_ / E)] (McLachlan and Martin, 1981). The value *f* is a factor that improves the accuracy of non-linear summation with a consideration of the capacitance of the muscle membrane and is set to 0.8 (Martin, 1976; McLachlan and Martin, 1981). E is the difference between the resting membrane potential (V_m_) and the reversal potential for ACh current, which is assumed as 0 mV (Magleby and Stevens, 1972a, b). The quantal content (the number of acetylcholine quanta released in response to a single nerve impulse) was calculated by the direct method using the formula Quantal Content = EPP’ / mEPP_normalized_ (Boyd and Martin, 1956; Wood and Slater, 2001). Rise time was defined as the time required for the response to increase from 10% to 90% of peak amplitude. Decay time was defined as the time for the response to decay from 100% to 50% (mEPPs) or from 90% to 10% (EPPs) of peak amplitude.

### Quantitative real-time PCR

RT-qPCR was carried out as previously described (Liu and Lin, 2023). Briefly, total RNAs were isolated from diaphragm muscles of 4 pairs of *Acta1^−/−^* and their littermate wildtype mice at P5 using TRI reagent (Molecular Research Center). First strand cDNAs were synthesized using iScript cDNA synthesis kit (BIO-RAD). Quantitative real time PCR was carried out by using iTaq Universal SYBR Green Supermix (BIO-RAD) on a QuantStudio 6 Pro Real-Time PCR System (ThermoFisher). Housekeeping gene GAPDH was used as an internal control. The relative expression levels of the genes of interest were normalized to the levels of GAPDH of the same sample by using DDCt method. The following primers were used for PCR amplification: (1) GAPDH primers, forward TCA ACG GCA CAG TCA AGG CCG AGA, reverse ATG ACC TTG CCC ACA GCC TTG GCA GC (Cossins et al., 2004); (2) AChR α subunit primers, forward CGT CTG GTG GCA AAG CT, reverse CCG CTC TCC ATG AAG TT; (3) AChR β subunit primers: forward CAA GGC ACC ATG CTC AGC CTC, reverse TCA GGA GCT ACG AGA GGT CAT; (4) AChR δ subunit primers, forward GTG ATC TGT GTC ATC GTA CT, reverse GCT TCT CAA ACA TGA GGT CA; (4) AChR ε subunit primers, forward AGA CCT ACA ATG CTG AGG AGG, reverse GGA TGA TGA GCG TAT AGA TGA; (5) AChR γ subunit primers, forward ACG GTT GTA TCT ACT GGC TG, reverse GAT CCA CTC AAT GGC TTG C (Tang and Goldman, 2006); (6) Rapsyn primers: forward ATATCGGGCCATGAGCCAGTAC, reverse TCACAACACTCCATGGCACTGC (Wang et al., 2008); (7) MuSK primers: forward CTCGTCCTCCCATTAATGTAAAAA, reverse TCCAGCTTCACCAGTTTGGAGTAA (Tang et al., 2009).

### Data analyses

Statistical analyses were performed using the number of animals (N) as the biological replicate. Data are presented as mean ± SEM. Two-tailed Student’s *t*-tests were used to compare mutant and control groups for endplate area, mEPP and EPP parameters, quantal content and mRNA levels. Paired t-tests were used for paired-pulse and repetitive stimulation analyses. Differences were considered statistically significant at *P* < 0.05.

## Author contributions

Yun Liu and Weichun Lin designed research; Yun Liu and Qiaohong Ye performed research; Yun Liu analyzed data; Yun Liu and Weichun Lin wrote the paper.

## Acknowledgements

We thank Dr. Kristen Nowak (The University of Western Australia) for providing heterozygous *Acta1*^+/−^mice originally generated by Dr. James Louis Lessard (Cincinnati Children’s Hospital Medical Center). We are grateful to Drs. Jane Johnson, Beverly Rothermel, and Shilpi Singh for their critical reading of earlier versions of the manuscript. This work was supported by grants from the National Institutes of Health/National Institute of Neurological Disorders and Stroke (R01 NS055028), the Edward Mallinckrodt, Jr. Foundation, and the Cain Foundation for Medical Research.

